# Secreted Proteases Control the Timing of Aggregative Community Formation in *Vibrio cholerae*

**DOI:** 10.1101/2021.05.25.445717

**Authors:** Matthew Jemielita, Ameya A. Mashruwala, Julie S. Valastyan, Ned S. Wingreen, Bonnie L. Bassler

## Abstract

Bacteria orchestrate collective behaviors using the cell-cell communication process called quorum sensing (QS). QS relies on the synthesis, release, and group-wide detection of small molecules called autoinducers. In *Vibrio cholerae*, a multicellular community aggregation program occurs in liquid, during stationary phase, and in the high-cell-density QS state. Here, we demonstrate that this aggregation program consists of two subprograms. In one subprogram, which we call void formation, structures form that contain few cells but provide a scaffold within which cells can embed. The other subprogram relies on flagellar machinery and enables cells to enter voids. A genetic screen for factors contributing to void formation, coupled with companion molecular analyses, showed that four extracellular proteases, Vca0812, Vca0813, HapA, and PrtV control the onset timing of both void formation and aggregation, and moreover, proteolytic activity is required. These proteases, or their downstream products, can be shared between void-producing and non-void-forming cells and can elicit aggregation in a normally non-aggregating *V. cholerae* strain. Employing multiple proteases to control void formation and aggregation timing could provide a redundant and irreversible path to commitment to this community lifestyle.

**Importance:** Bacteria can work as collectives to form multicellular communities. *Vibrio cholerae*, the bacterium that causes the disease cholera in humans, forms aggregated communities in liquid. Aggregate formation relies on a chemical communication process called quorum sensing. Here we show that, beyond overarching control by quorum sensing, there are two aggregation subprograms. One subprogram, which we call void formation, creates a scaffold within which cells can embed. The second subprogram, which allows bacteria to enter the scaffold, requires motility. We discovered that four extracellular proteases control the timing of both void formation and aggregation. We argue that, by using redundant proteases, *V. cholerae* ensures the reliable execution of this community formation process. These findings may provide insight into how *V. cholerae* successfully alternates between the marine environment and the human host: transitions that are central to the spread of the disease cholera.

## Introduction

Bacteria often form multicellular communities. In *Vibrio cholerae*, the pathogen responsible for the disease cholera, multicellular community formation is controlled by the bacterial cell-cell communication process called quorum sensing (QS). QS relies on extracellular signal molecules called autoinducers. Autoinducers are detected by the population, facilitating collective behaviors.

A simplified schematic of the *V. cholerae* QS circuit showing components germane to the present work is provided in Fig. 1 (1). In brief, when autoinducer concentration is low, the autoinducer receptors act as kinases that shuttle phosphate to the master response regulator LuxO. LuxO-P drives AphA production and represses HapR production. AphA and HapR are, respectively, the master low-cell-density (LCD) and high-cell-density (HCD) QS transcriptional regulators. When autoinducer concentration is high, the receptors act as phosphatases that promote the removal of phosphate from LuxO. Dephosphorylated LuxO is inactive so AphA is no longer made and HapR production is no longer repressed. HapR activates expression of genes in the HCD QS regulon.

**Figure 1:**
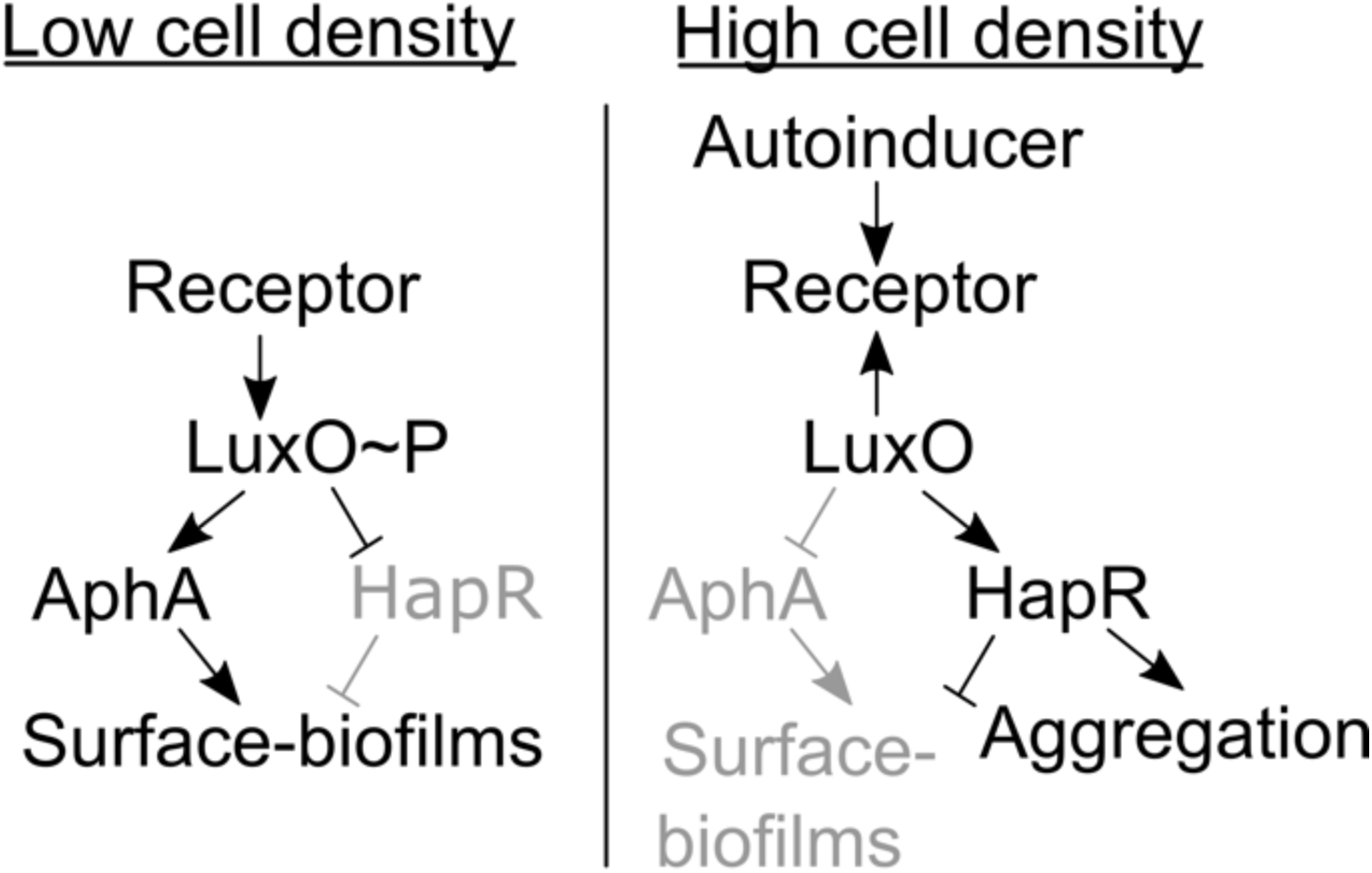
Simplified *V. cholerae* QS circuit. See text for details.

In the LCD QS state, *V. cholerae* forms surface-biofilms: bacterial communities bound to each other and to surfaces by an extracellular matrix. In the HCD QS state, *V. cholerae* disperses from surface-biofilm communities and reenters the planktonic, individual-cell lifestyle (2, 3). To carry out the present work, we used *V. cholerae* strains locked in the LCD and HCD QS modes. The strain locked in the LCD QS mode carries the *luxO D61E* mutation, encoding a LuxO phosphomimetic. The strain locked in the HCD QS mode carries the *luxO D61A* mutation, encoding a LuxO variant incapable of being phosphorylated (4).

We previously reported a liquid-based aggregative community formation program in *V. cholerae* (5). This program is launched in the HCD QS state and is positively regulated by HapR (Fig. 1) (5). By contrast, *V. cholerae* surface-biofilm formation occurs at LCD. Formation of surface biofilms requires *Vibrio* polysaccharide (VPS) production (6–8). The aggregation program does not require VPS. Maturation of surface biofilms depends on cell division and takes many hours. From inception to completion, the *V. cholerae* aggregation program takes at most 30 min, precluding a cell-division-driven mechanism (5). These differences suggest that distinct mechanisms underlie surface-biofilm formation and aggregation. Previously, we performed a genetic screen to identify factors promoting aggregation. That screen revealed that motility and stress response genes, among others, are required (5).

Here, we further investigate the *V. cholerae* aggregation program. To ensure that we study the process independently of surface-biofilm formation, unless otherwise noted, all strains lack *vpsL*, a gene essential for VPS production (9). We show that the *V. cholerae* aggregation program involves structures that we call voids. Voids can form in *V. cholerae* strains in which the flagellar machinery is disrupted. Voids contain few cells. Additionally, if all cells are removed from a culture shortly before the onset of void formation, voids still form in their absence. Thus, voids presumably provide a scaffold within which cells embed to form aggregative communities. We demonstrate that the onset timing of void formation, and therefore, the onset timing of aggregation, are controlled by four extracellular proteases: Vca0812, Vca0813, HapA, and PrtV. Using site directed mutagenesis of the Vca0812 protease as a test case, we show that proteolytic activity is necessary for proper void/aggregation onset timing. The four proteases can be shared among void-forming and non-void-forming strains. Indeed, protease-deficient *V. cholerae* strains that exhibit delayed void formation and aggregation timing were restored to wildtype aggregation timing by incubation with protease-harboring strains. We propose a model in which proteases cleave a substrate(s), converting a precursor into a product that promotes void formation/aggregation or one that loses the ability to repress void formation/aggregation. Possibly, the involvement of four redundant proteases ensures the rapid and reliable execution of this *V. cholerae* multicellular program.

## Results

### The *V. cholerae* aggregation program relies on a void formation subprogram

We previously performed a transposon mutagenesis screen in the Δ*vpsL* HCD-locked *V. cholerae* strain to identify components required for aggregation (5). This screen revealed that a *flgC*::Tn*5* Δ*vpsL* HCD-locked mutant did not participate in aggregative community formation in contrast to its Δ*vpsL* HCD-locked parent strain (Fig. 2A), and rather formed structures in liquid with few embedded cells (Fig. 2B). *flgC* encodes a flagellar basal body rod protein (11). We call the structures made by this mutant ‘voids’. Voids can be visualized using India ink negative staining (12) (Fig. 2A-C). We validated the phenotype of the *flgC*::Tn*5* Δ*vpsL* HCD-locked mutant by constructing an in-frame deletion of *flgC* to generate the Δ*flgC* Δ*vpsL* HCD-locked strain (Fig. 2C). Indeed, relative to the Δ*vpsL* HCD-locked parent strain, both the strain with the transposon insertion and the strain with the deletion in *flgC* formed voids that largely lacked embedded cells (Fig. 2D). This finding suggested that void formation is an aspect of the overall aggregation program.

**Figure 2:**
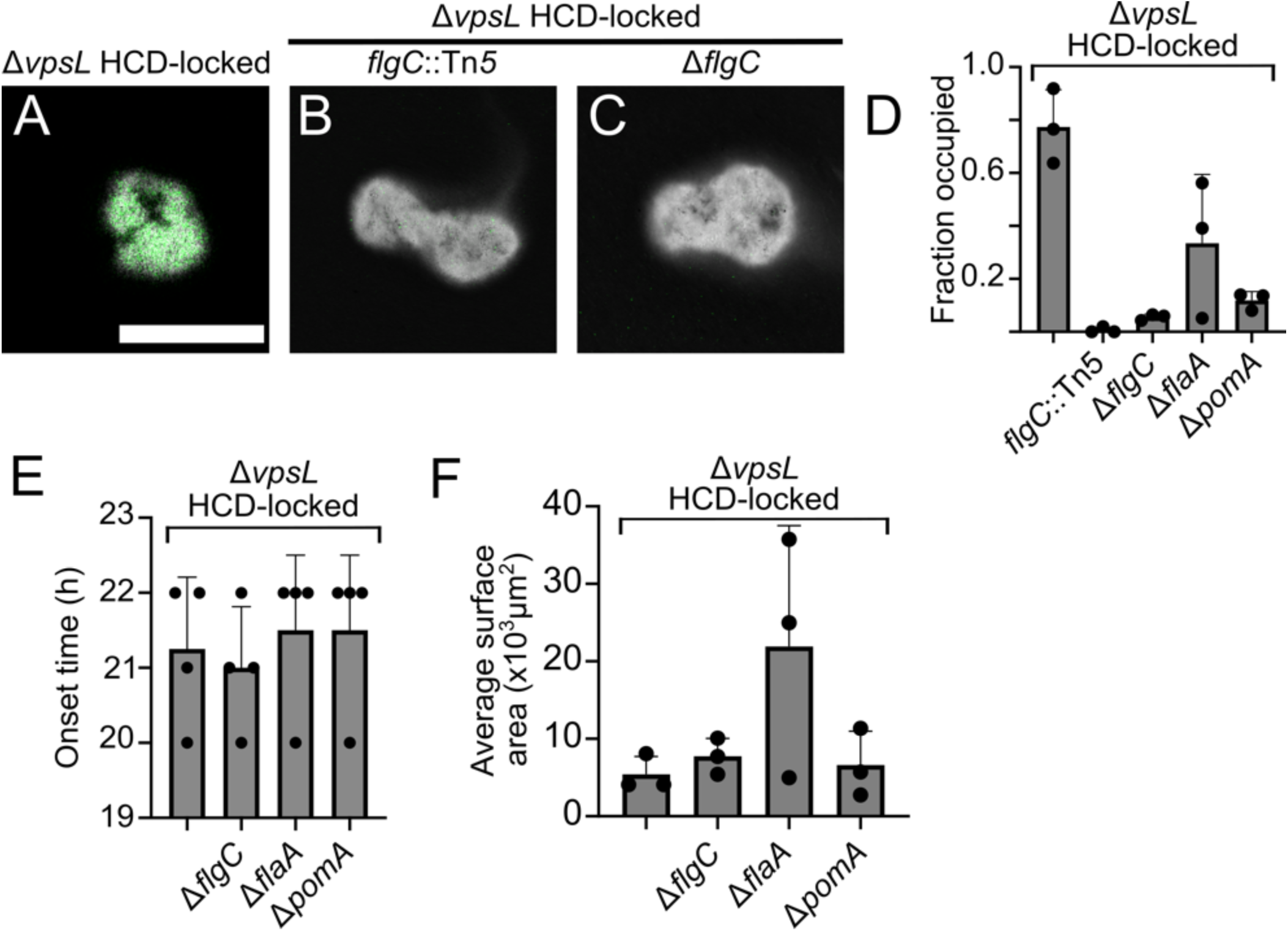
The *V. cholerae* aggregation program contains a void formation sub-program. Representative cross-sectional images of the designated (A) aggregating (B, C) void-forming strains. (A-C) treated with India ink counterstain (gray; inverted lookup table). Green: SYTO-9 nucleic acid stain. Scale bar: 100 µm. Magnification: 63x. (D) Quantitation of average occupancy of voids or aggregates at T = 22 h. Error bars denote mean ± standard deviation (SD), N = 3 biological replicates. (E) Quantitation of onset time of designated strains relative to the Δ*vpsL* HCD-locked strain. Error bars denote mean ± SD, N = 4 biological replicates. (F) Quantitation of average cross-sectional surface area for the strains in (E). Error bars denote mean ± SD, N = 3 biological replicates. (B-F) All strains except the *flgC*::Tn*5* Δ*vpsL* HCD-locked mutant constitutively expressed *mKO* from the chromosome.

To study features of void formation and aggregation, we focused on two readily assayable phenotypes: onset timing and structure size. Every 1 h, we imaged the Δ*flgC* Δ*vpsL* HCD-locked and Δ*vpsL* HCD-locked strains each containing a chromosomally integrated fluorescent *mKO* reporter. Void formation and aggregation, respectively, occurred with similar timing in the two strains (Fig. 2E) and the voids and aggregates were comparable in size (Fig. 2F).

We tested whether the inability of cells to enter voids was specific to the Δ*flgC* Δ*vpsL* HCD-locked mutant or, alternatively, whether possessing a flagellum and/or being motile was required. For this analysis, we examined two additional strains: Δ*flaA* Δ*vpsL* HCD-locked and Δ*pomA* Δ*vpsL* HCD-locked. *flaA* encodes an essential flagellum subunit. Thus, Δ*flaA* mutants have no flagella. *pomA* encodes the stator complex of the flagellar motor and is required for flagellar rotation (13). Thus, Δ*pomA* mutants have flagella, but they do not rotate. Void formation timing was the same for all of these strains (Fig. 2E). The voids formed by the Δ*flaA* Δ*vpsL* HCD-locked strain harbored more embedded cells than voids formed by the Δ*pomA* Δ*vpsL* HCD-locked and Δ*flgC* Δ*vpsL* HCD-locked strains, but harbored fewer cells than the voids formed by the Δ*vpsL* HCD-locked parent (Fig. 2D). The Δ*flaA* Δ*vpsL* HCD-locked strain also made larger structures than those made by the other strains (Fig. 2F). We conclude that *V. cholerae* void formation does not require flagella or motility. However, flagella capable of rotation are necessary for cells to enter voids and form aggregates. We do not yet know if motility per se is required, or rather, if flagellar rotation plays a regulatory or adhesive role in aggregate formation. Additionally, there appears to be FlaA-mediated regulation of void size and cell entry (Fig. 2D, F). Mutation of *flaA* caused pleotropic effects in other studied *V. cholerae* strains (14). We do not investigate the FlaA-mediated effect further in the current work.

Based on the above findings, we propose that the aggregation program consists of two subprograms: one subprogram is void formation, in which structures are made within which cells can embed, and the other subprogram uses the flagellar machinery to facilitate cell entry into voids to form aggregates. Here, we used the Δ*flgC* Δ*vpsL* HCD-locked strain as our model strain that only engages in the void formation subprogram. We call the Δ*flgC* Δ*vpsL* HCD-locked strain the “Void^+^” strain. We call the Δ*vpsL* HCD-locked strain that undergoes the full aggregation program the “Aggregate^+^” strain.

### A genetic screen identifies *V. cholerae* factors that promote void formation

We reasoned that we could identify factors contributing to void formation by mutagenizing the Void^+^ strain and screening for defects in this process. To accomplish this, we needed to rapidly distinguish void-forming from non-void-forming mutants. The Aggregate^+^ and the Void^+^ strains can be differentiated from the non-aggregating Δ*vpsL* LCD-locked strain based on colony morphology: on agar plates, Void^+^ and Aggregate^+^ colonies are opaque, while non-aggregating Δ*vpsL* LCD-locked colonies are translucent. We mutagenized the Void^+^ strain with Tn*5* and screened ∼65,000 colonies for those that obtained translucent phenotypes, suspecting that some could be non-void-forming mutants. We used a secondary microscopy screen to assess void formation. Our strategy yielded 92 putative candidates with mutations mapping to 25 loci (Table S1). The mutated genes encoded the regulatory proteins RpoS, CyaA, HapR, VarS, and SspA, and genes encoding biosynthetic and metabolic enzymes, among others. Here, we focus on one identified operon, *vca0812-vca0813*, that encodes two putative extracellular proteases: Vca0812 and Vca0813 (15, 16). *vca0812* and *vca081*3 are occasionally called *lap* and *lapX* (16, 17). Below, we first study Vca0812 and Vca0813 in the Void^+^ background to determine how they contribute to void formation, and second, we assess their roles in the Aggregate^+^ background to define their effects on aggregation.

### The extracellular proteases Vca0812 and Vca0813 control the onset timing of *V. cholerae* void formation and aggregation

To study the roles of Vca0812 and Vca0813 in void formation, we deleted *vca0812* and *vca0813* individually and together in the Void^+^ parent strain. We assayed the mutants for void formation capability relative to the parent Void^+^ strain. The strains lacking either or both proteases showed ∼3.5 h delays in void formation onset (Fig. 3A), however, ultimately, they formed voids. We conclude that Vca0812 and Vca08123 control void formation onset timing.

**Figure 3:**
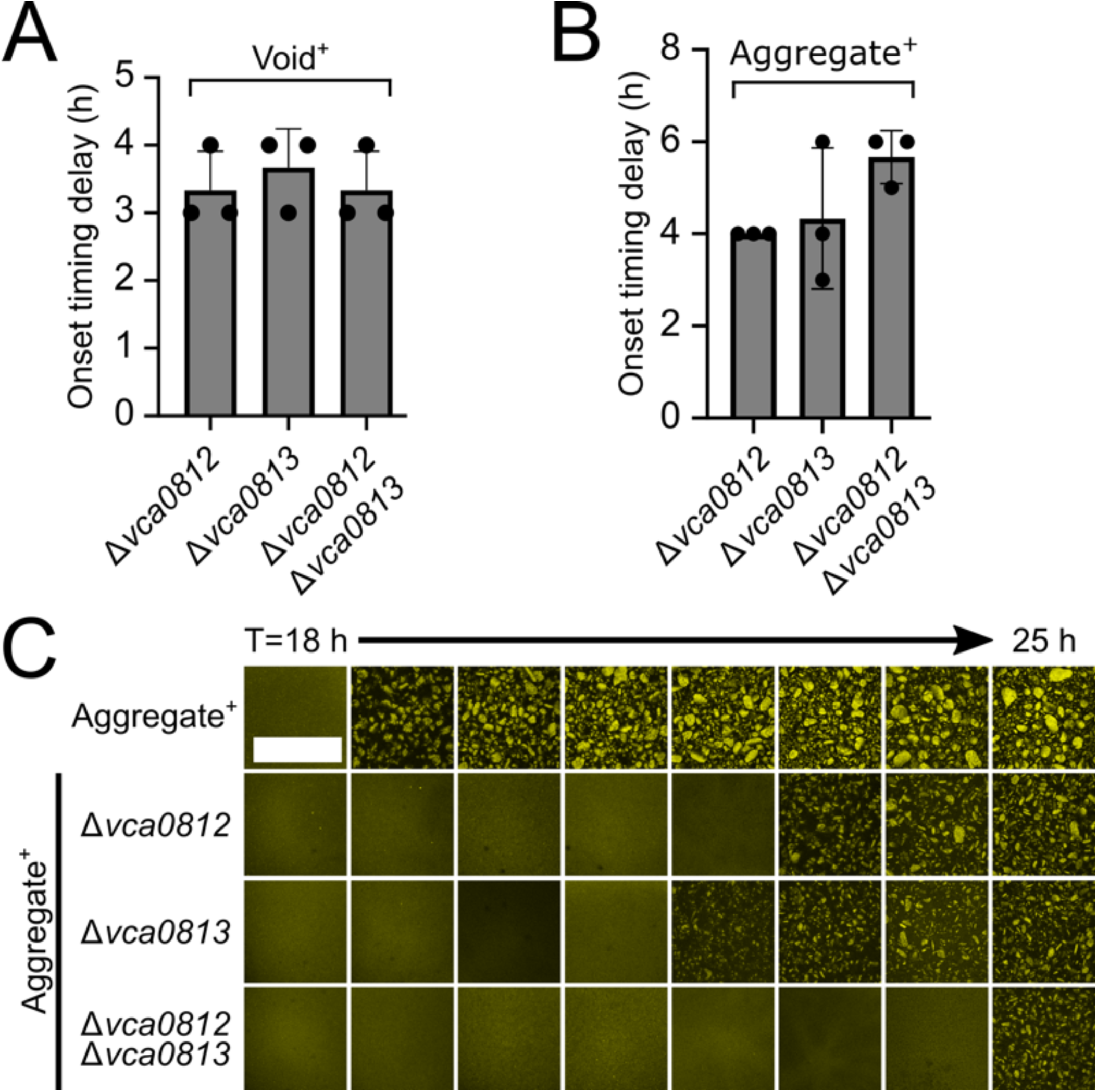
A genetic screen reveals *vca0812* and *vca0813* as regulators of *V. cholerae* void formation onset timing. (A) Quantitation of void formation onset timing delay for the designated strains relative to that of the Void^+^ strain. (B) Quantitation of aggregation onset timing delay for the designated strains relative to that of the Aggregate^+^ strain. (A, B) Error bars denote mean ± SD, N = 3 biological replicates. (C) Representative cross-sections through the designated strains imaged every 1 h. Scale bar: 1000 µm. Magnification: 10x. (A-C) All strains constitutively expressed *mKO* from the chromosome.

To examine whether Vca0812 and Vca0813 also affect aggregation onset timing, we constructed the same deletions in the Aggregate^+^ background. Deletion of *vca0812* and/or *vca0813* resulted in aggregation onset delays of ∼4 h or more compared to the parent Aggregate^+^ strain (Fig. 3B, C). There were no growth defects in any of the strains containing the *vca0812* and *vca0813* deletions (Fig. S1A). To verify that Vca0812 and Vca0813 drive aggregation onset timing, we complemented the Δ*vca0812* Aggregate^+^ and Δ*vca0813* Aggregate^+^ strains with, respectively, *vca0812* and *vca0813* under the native promoter from an ectopic chromosomal locus. In both cases, wildtype (WT) aggregation timing was restored (Fig. S2). Finally, GbpA, a *V. cholerae* adhesin (18, 19) that is encoded by the *vca0811* gene and located immediately adjacent to the *vca0812*-*vca0813* operon does not contribute to void formation or aggregation (Fig. S3). We conclude that Vca0812 and Vca0813 control void formation onset timing, which, consequently, affects aggregation onset timing. However, other components must also participate in void formation and in the aggregation program because in the absence of these two proteins voids still form and aggregation occurs, albeit with delayed timing.

### The secreted proteases HapA and PrtV also contribute to void and aggregate onset timing

Given that the extracellular proteases Vca0812 and Vca0813 control both void and aggregation onset timing, and their elimination did not fully abrogate either process, other *V. cholerae* extracellular proteases were candidates for program control. The *V. cholerae* genome is predicted to encode genes specifying seven additional extracellular proteases: VesA, VesB, VesC, IvaP, TagA, PrtV, and HapA (16, 17, 20–22). We deleted the genes encoding six of these proteases in the Void^+^ strain background. Our attempts to construct an in-frame deletion of *tagA* were unsuccessful, so we engineered a *tagA*::Kan^R^ strain. Deletion of *hapA* and *prtV* caused void formation onset timing delays while deletion of *vesA*, *vesB*, *vesC*, or *ivaP*, and transposon insertion into *tagA*, did not (Fig. 4A). Compared to the Void^+^ parent, the Void^+^ strain lacking all four relevant protease-encoding genes (Δ*vca0812* Δ*vca0813* Δ*hapA* Δ*prtV*; designated Δ*4* Void^+^) exhibited severely delayed void formation onset timing, ∼14 h, i.e., longer than that of any mutant lacking any single protease (Fig. 4A). Using the Aggregate^+^ strain as the parent, we showed that HapA and PrtV played analogous roles in delaying aggregate onset formation, and, again, simultaneous elimination of all four relevant proteases caused the most severe delay (Fig. 4B). Proper timing, or nearly proper timing in the case of PrtV, was restored following complementation with the cloned genes (Fig. S2). Again, there were no growth defects (Fig. S1B). We conclude that, in addition to Vca0812 and Vca0813, the HapA and PrtV extracellular proteases contribute to the control of void and aggregate onset timing. We also tested whether elimination of these four proteases delayed aggregation in an otherwise WT strain (i.e., in a strain harboring an intact QS circuit, that produces VPS, and possesses a functional flagella). Indeed, WT *V. cholerae* in which the *vca0812*, *vca0813*, *hapA*, and *prtV* genes had been deleted exhibited a severe onset delay in aggregation (Fig. 4B).

**Figure 4:**
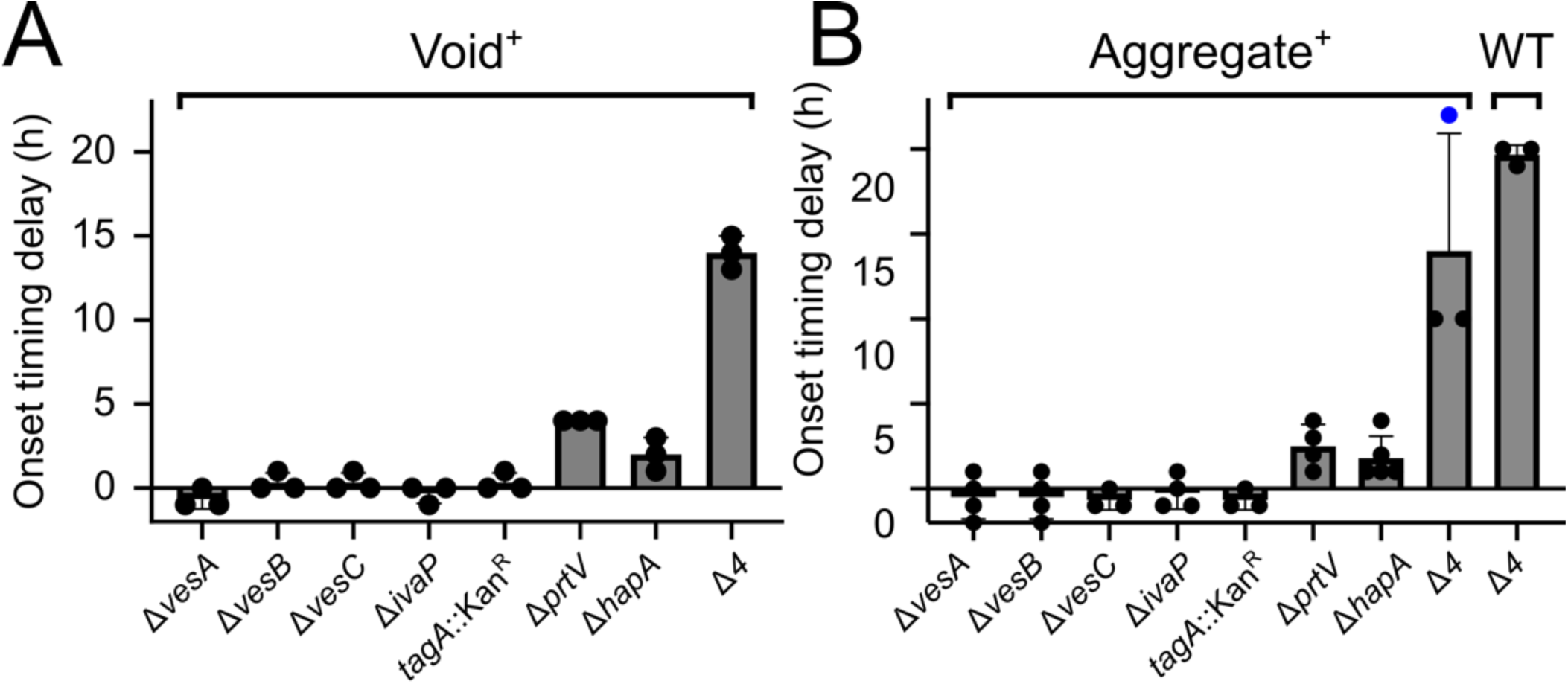
HapA and PrtV regulate *V. cholerae* void formation and aggregation onset timing. (A) Quantitation of void formation onset timing delay of the designated strains relative to that of the Void^+^ strain. (B) Quantitation of aggregation onset timing delay of the designated strains relative to that of the Aggregate^+^ strain. The blue circle indicates a sample that did not exhibit aggregate formation at the assayed time point. (A, B) Error bars denote mean ± SD, N = 3 biological replicates. The Void^+^ and Aggregate^+^ strains were assayed until T = 32 h. The WT strain was assayed until T = 26 h. All strains were assayed again at T = 42 h if no aggregation/void formation had occurred by the earlier timepoint. All strains constitutively expressed *mKO* from the chromosome.

### Proteolytic activity is required for control of void and aggregation onset timing

A model capturing our above findings is that Vca0812, Vca0813, PrtV, and/or HapA proteolytic activity is required for void formation and aggregation to commence with WT timing. We predict that these proteases cleave a substrate. Cleavage either converts a precursor into an active form required for void formation and aggregation, or alternatively, cleavage inactivates an inhibitor that suppresses void formation and aggregation. To test the necessity for proteolytic activity, we employed a mutant defective in proteolytic activity and, independently, we chemically perturbed protease activity. Regarding the catalytic mutant, we focused on Vca0812 and Vca0813 because they exert the strongest effects on void formation and aggregation onset timing (see Figs. 3 and 4). Crystal structures of Vca0812 and Vca0813 do not exist, so we used I-TASSER to thread Vca0812 and Vca0813 onto homologous proteins from the Protein Data Bank (23). This strategy yielded a predicted catalytic triad for Vc0812: His191, Asp236, and Ser319. Parallel efforts with Vca0813 were unsuccessful. To test the requirement for catalysis in void formation, we introduced pBAD expressing either Vca0812 or Vca0812 H191N into the Δ*4* Void^+^ strain. Zymography showed a proteolytically-active band in the strain carrying *vca0812* that is absent from the strain carrying *vca0812* H191N and the strain harboring the empty vector control (Fig. 5A). Analogous results were obtained with the Δ*4* Aggregate^+^ strain (Fig. 5B). Western blotting verified that the amounts of Vca0812 and Vca0812 H191N were similar in the Δ*4* Void^+^ (Fig. S4A) and Δ*4* Aggregate^+^ (Fig. S4B) strains, although the mutant protein exhibited somewhat higher susceptibility to degradation in the Δ*4* Aggregate^+^ strain (Fig. S4B). We conclude that Vca0812 has proteolytic activity and that Vca0812 H191N is defective for catalysis. Void formation onset timing in the *vca0812 H191N* Void^+^ strain was similar to that of the Δ*vca0812* Void^+^ strain, i.e., they respectively had an ∼3 and 3.3 h delay, relative to the parent Void^+^ strain (Fig. 5C). In the case of aggregation, the timing delay of the *vca0812 H191N* Aggregate^+^ strain was similar to that of the Δ*vca0812* Aggregate^+^ strain (2 h for both, Fig. 5D). We conclude that Vca0812 proteolytic activity is required for proper onset timing of void formation and aggregation.

**Figure 5:**
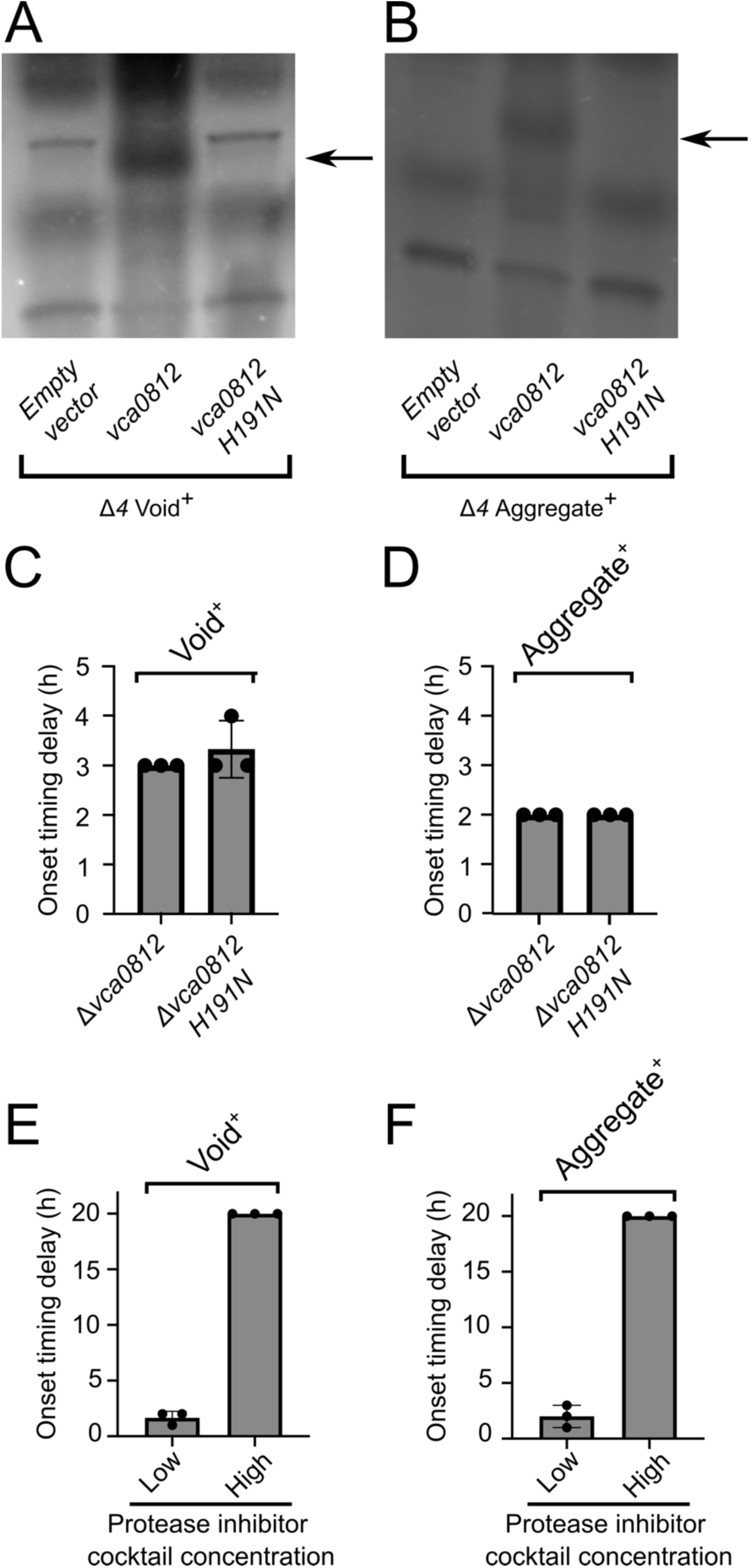
Vca0812 proteolytic activity is required for proper *V. cholerae* void and aggregation onset timing. (A) Zymographic analyses of proteolytic activity in cell-free culture fluids from the Δ*4* Void^+^ strain, expressing the indicated genes from pEVS-pBAD. (B) As in (A) for the Δ*4* Aggregate^+^ strain. (A, B) Arrows indicate the regions corresponding to Vca0812 activity. Gels were cropped to show the relevant regions and the look-up table was inverted. The presence of a band indicates proteolytic activity. (C) Quantitation of void formation onset timing delay for the designated strains relative to that of the Void^+^ strain. (D) Quantitation of aggregation onset timing delay for the designated strains relative to that of the Aggregate^+^ strain. (E) Quantitation of void formation onset timing delay for the Void^+^ strain to which a low or high concentration of a protease inhibitor cocktail was administered at T = 0 h, relative to the onset time of the Void^+^ strain. (F) Quantitation of aggregation onset timing delay of the Aggregate^+^ strain to which a low or high concentration of a protease inhibitor cocktail was added at T = 0 h, relative to the onset time of the Aggregate^+^ strain. (E, F) Samples were assayed from T = 22 h until T = 26 h, and then again at T = 42 h if no void formation or aggregation had occurred by T = 26 h. The low protease cocktail concentration was 1 tablet of Roche cOmplete protease inhibitor cocktail per 100 mL and the high protease cocktail concentration was 1 tablet per 20 mL. (C-F) All strains constitutively expressed *mKO* from the chromosome.

In an independent test for the requirement for proteolytic activity in void formation and aggregation, we used a broad-spectrum protease inhibitor cocktail (see Methods). We added it to the Void^+^ and Aggregate^+^ strains at T = 0 h. Low and high inhibitor concentrations drove ∼2 and >4 h delays, respectively, in void formation onset timing in the Void^+^ strain (Fig. 5E). In the Aggregate^+^ strain, ∼1.6 and >4 h aggregation delays occurred following, respectively, low and high inhibitor treatments (Fig. 5F). Only modest growth rate reductions occurred when the protease inhibitor was present (Fig. S1C), suggesting that changes in growth rate are not responsible for differences in void formation onset timing. We conclude that proteolytic activity contributes to the proper onset timing of void and aggregate formation.

### Extracellular proteases can be shared during *V. cholerae* void and aggregate formation

Our finding that extracellular proteases are involved in void and aggregate formation suggested the possibility that, in the absence of cells, components in cell-free fluids might be sufficient to drive the process. To test this idea, immediately prior to void formation onset, we removed the Aggregate^+^ cells from their growth medium and allowed the conditioned medium to continue to incubate (Fig. 6A). Structures resembling voids (Fig. 2B, C) spontaneously formed (Fig. 6B). We call these structures “cell-free voids”. When such conditioned medium was prepared from Void^+^ strains from which we had deleted *vca0812*, *vca0813*, *prtV*, *hapA*, or all four of these genes (Δ*4*), the preparations were incapable (Δ*vca0812*, Δ*vca0813,* Δ*prtV*, Δ*4*) or severely defective (Δ*hapA*) in promoting cell-free void formation (Fig. 6C). By contrast, conditioned medium prepared from Void^+^ strains from which we had deleted *vesA*, *vesB*, *vesC*, or *ivaP*, or inactivated *tagA* by transposon insertion, (i.e., genes encoding the proteases for which we found no role in void and aggregate formation) formed cell-free voids with the same onset time as those formed in the conditioned medium from the parent Void^+^ strain (Fig. 6C). Parallel results were obtained for the Aggregate^+^ strain, with one difference: conditioned medium from strains lacking *hapA* or *prtV* exhibited heterogeneity in onset timing delay (Fig. 6D). Combined, these data argue that the four identified extracellular proteases function as factors to promote cell-free void formation.

**Figure 6:**
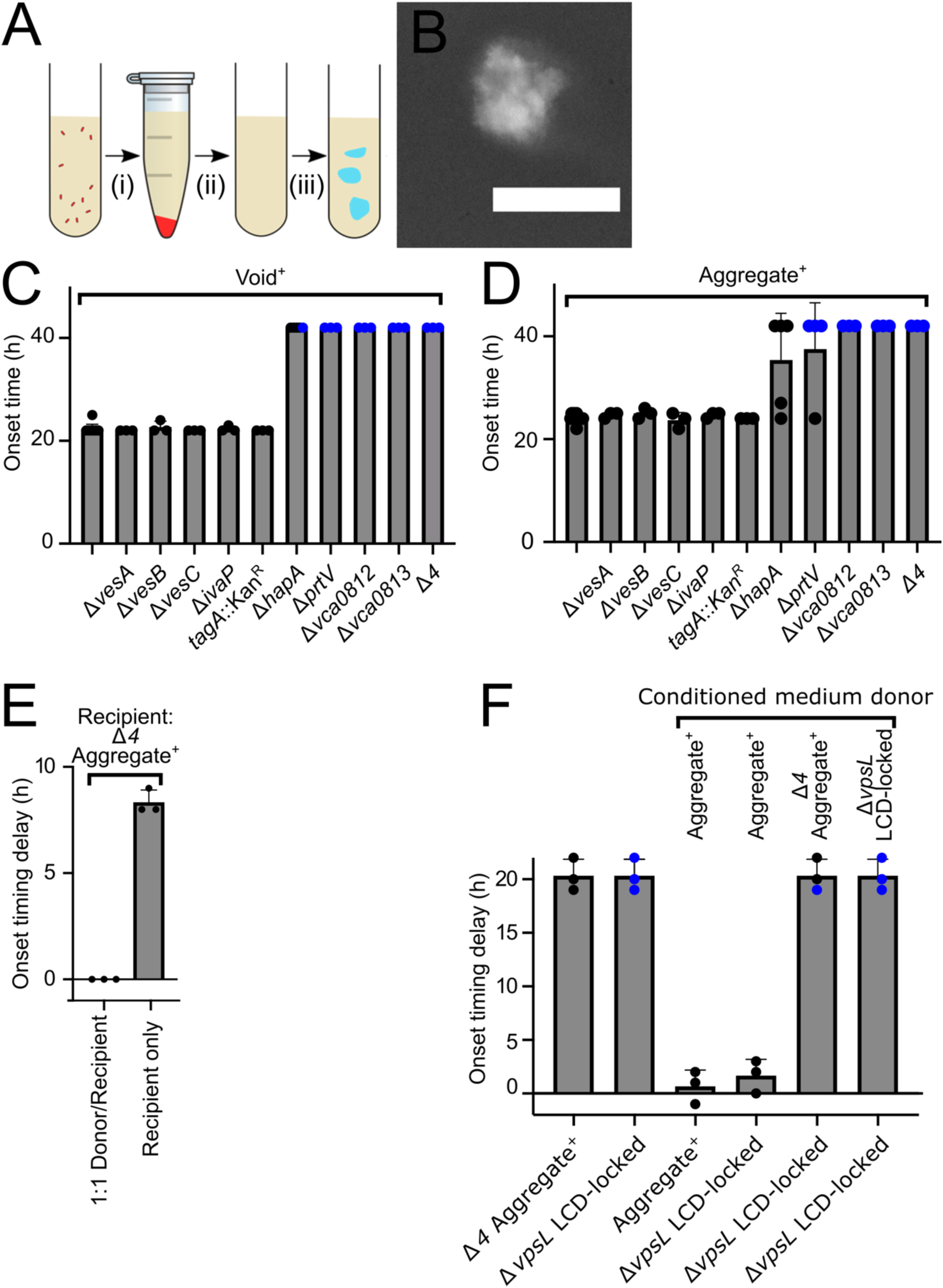
Cell-free conditioned medium forms voids and the void-promoting activity can be cross-fed to recipient cells. (A) Protocol for cell-free void isolation. Cells were removed via centrifugation (i) and the conditioned medium was filter sterilized (ii). This preparation was further incubated and cell-free voids formed (iii). (B) Image of a cell-free void (protocol step iii in (A)). Gray: India ink counterstain (inverted look-up table). Scale bar: 100 µm. Magnification: 63x. (C) Onset time of cell-free void formation for the designated strains in the Void^+^ strain background. (D) Onset time of cell-free void formation for the designated strains in the Aggregate^+^ strain background. (C, D) Error bars are mean ± SD, N = 3 biological replicates. Conditioned medium generated at T = 18 h. Strains assayed from T = 22 h until T = 26 h, and then again at T = 42 h if no cell-free void formation had occurred by T = 26 h. (E) Aggregation onset delay for a co-culture of the Aggregate^+^ (donor) and Δ*4* Aggregate^+^ (recipient) strain and the Δ*4* Aggregate^+^ strain alone relative to the onset timing of the Aggregate^+^ strain. The Aggregate^+^ and the Δ*4* Aggregate^+^ strain constitutively expressed *mKate2* and *mKO* respectively, from the chromosome. Error bars are mean ± SD, N = 3 biological replicates. (F) Aggregation onset delay for the designated strains that had been supplied the indicated conditioned medium at T = 18 h relative to the onset timing of the Aggregate^+^ strain to which no conditioned medium was supplied. Samples were assayed from T = 22 h until T = 26 h, and then again at T = 42 h if no void formation or aggregation had occurred by T = 26 h. (C, D, F) Blue circles indicate samples that did not exhibit void formation or aggregation at the assayed time point. Strains in B, C, D, F constitutively expressed *mKO* from the chromosome.

Bacterial exoproducts are often shared among cells within communities (24). To explore whether this is the case for the proteases controlling void and aggregation formation onset timing, we co-cultured protease-producing strains with strains lacking the proteases and assayed whether the non-producing strains regained WT aggregation timing. Specifically, we combined equal amounts of two strains at T = 0 h. The donor strain was the Aggregate^+^ strain, which carries all the extracellular proteases required for proper void formation and aggregation timing. The recipient strain was the Aggregate^+^ strain lacking the four proteases that control void formation and aggregation onset timing (Δ*4* Aggregate^+^ strain). The donor and recipient strain were, respectively, labeled with *mKate2* and *mKO*, which enabled us to track the cells of each strain. Co-culture allowed the defective recipient strain to regain the shorter, WT aggregation onset timing (Fig. 6E). Importantly, the aggregates that formed in the co-culture contained both Aggregate^+^ donor and protease-deficient recipient cells (Video S1). Co-culture of the Aggregate^+^ donor with recipient strains lacking only one of the key proteases (i.e., Δ*vca0812*, Δ*vca0813*, Δ*prtV*, or Δ*hapA*) also shortened the duration of the recipients’ delay in aggregation onset timing (Fig. S5). We conclude that the activity of these proteases, and/or the components that are processed by them, can be shared among cells.

We also examined whether providing the proteases in fluids lacking donor cells could rescue a mutant incapable of aggregation. To do this, we generated cell-free conditioned medium from the Aggregate^+^ strain and the Δ*4* Aggregate^+^ strain. We supplied these preparations to the non-aggregating Δ*vpsL* LCD-locked strain at T= 18 h. Conditioned medium from the Aggregate^+^ strain elicited aggregation, while conditioned medium from the Δ*4* Aggregate^+^ strain did not (Fig. 6F). We conclude that proteases present in culture fluids are sufficient to restore aggregation to a non-aggregating recipient strain. At present, we do not know whether the shared component is the protease itself, the product of protease digestion, or both.

## Discussion

Aggregative community formation is a program of *V. cholerae* multicellularity that occurs rapidly during stationary phase and when cells are in the HCD QS state (5). We propose that the aggregation program can be divided into two subprograms: One is responsible for void formation and the other, mediated by the flagellar machinery, facilitates cell entry into voids. In this work, we demonstrate that four extracellular proteases, Vca0812, Vca0813, HapA, and PrtV, control the onset timing of void formation and thus, aggregation. *V. cholerae* is predicted to possess nine total extracellular proteases (16, 17, 20–22). The four proteases for which we identified roles are reported to be the most active extracellular proteases during stationary phase (17). Moreover, QS activates the expression of the genes encoding them at HCD (17), consistent with the timing we discovered for aggregation onset (Fig. 3 and 4). Roles in controlling surface biofilm formation have been ascribed to HapA and PrtV (25, 26), but to our knowledge, no such roles have previously been assigned to Vca0812 or Vca0813.

As a test case for the involvement of these proteases, we demonstrated that Vca0812 proteolytic activity is required for proper void formation and aggregation program onset timing. While untested, we anticipate that the other three proteases likewise harbor proteolytic capability that contribute to program timing. We note that these proteases could additionally promote void formation and aggregation in a proteolysis-independent manner, for example, by acting as adhesins needed to assemble voids. The proteases are shared among community members as demonstrated by our finding that the conditioned medium from the protease proficient Aggregate^+^ strain can elicit aggregation in the normally non-aggregating Δ*vpsL* LCD-locked strain (Fig. 6F). Hundreds of genes are regulated differently in the Δ*vpsL* LCD-locked strain and the Aggregate^+^ strain since the entire QS regulon is in the LCD mode in the former strain and in the HCD mode in the latter strain. Thus, apparently, the crucial QS role in void and aggregate formation onset timing is the proper regulation of production of these extracellular proteases and/or regulation of their substrates. We propose that the identified extracellular proteases act on a target substrate(s) and cleave it into a product that fosters void formation and aggregation. Alternatively, cleavage could inactivate an inhibitor that suppresses void formation and aggregation. Substrate identification may be achieved by exploiting the catalytically defective Vca0812 H191N mutant, followed by substrate trapping, purification, and mass spectrometry analysis (27).

Program control by multiple proteases could yield benefits not achievable via the involvement of only a single protease. For example, redundancy could facilitate program fidelity by buffering against fluctuations in protease abundance or required cofactors (28). Additionally, two defining features of proteolytic regulation are that such processes are rapid relative to translational or transcriptional control and, furthermore, changes are irreversible (29). Proteolysis speed may underpin the rapid formation of aggregates (5) and the irreversibility of proteolytic cleavage could enable program commitment once the collective decision to undergo aggregation has been made (30). Again, having multiple proteases involved could ensure robustness or underpin a bet-hedging strategy (31) that protects aggregation program execution from competitors that can disable some, but not all, of the proteases involved in driving void and aggregate formation. Previously, we proposed that *V. cholerae* uses the aggregation formation program to rapidly assemble communities during infection and/or to enhance the successful dispersal from the human host back into the marine environment (5). If so, employing multiple, redundant proteases could ensure the reliable launch of the program in an environment as dynamic and complex as the human intestine.

In *V. cholerae* and other bacteria, proteolysis-dependent adhesion systems exist (25, 32–34). In some cases, protease activity reduces biofilm formation capacity. For example, in *Pseudomonas aeruginosa*, the periplasmic protease LapG cleaves the adhesin CdrA, releasing CdrA from the cell surface, the consequence of which is a reduction in biofilm formation (34). By contrast, proteolytic processing can also be essential for biofilm maturation. In *V. cholerae*, cleavage of RbmA by the extracellular protease HapA, PrtV, or IvaP, is important for the development of the WT biofilm architecture (35). Beyond prokaryotes, the void formation process has parallels to mechanisms for mammalian blood coagulation (36). When the vascular system sustains injury, a series of proteolytic events is initiated that converts fibrinogen, a soluble protein in blood, into fibrin, which rapidly polymerizes to form a clot. Our evidence suggests that void formation may employ an analogous strategy to rapidly form structures in liquid. In the case of void formation, we suspect that the putative protease substrate is extracellular, because cell-free voids form in conditioned medium lacking cells (Fig. 6B-D). This substrate may undergo auto-processing since elimination of all four proteases that, individually, alter onset timing does not fully eliminate void formation and aggregation. Alternatively, another extracellular protease, that we did not identify, could exist and fulfill this function.

Finally, formation of voids could occur by a phase transition mechanism. Proteolytic degradation of a substrate could convert it into a form that preferentially adheres to itself or to another component present in the cell-free fluids because of, for example, changes in charge or hydrophobicity. Because the void formation process is proteolytically driven, it is likely irreversible and the concentration of the proteolyzed product should increase over time until it achieves a level that allows the system to lower its free energy by spontaneously demixing. This process is called spinodal decomposition (37). Key features of spinodal decomposition are that the process is driven by local concentration fluctuations and it occurs spontaneously without the need to overcome an energetic barrier, as is the case with, for example, nucleation-driven phase separation. One difference between void formation and aggregation and classical theories of spinodal decomposition is that, in the latter case, demixing is initiated at a characteristic length scale, but further coarsens over time (37). By contrast, aggregates eventually stop enlarging (5). In future work, we aim to quantitatively study void formation to determine whether spinodal decomposition provides an appropriate framework for their understanding. In summary, our work demonstrates that extracellular proteases play a key role in controlling the onset of the *V. cholerae* aggregative community formation program.

## Materials and Methods

### Reagents and bacterial cultures

The parent strain was *V. cholerae* O1 El Tor biotype C6706str2 (38). When antibiotics were required, they were used at the following concentrations: ampicillin, 100 mg/L; kanamycin 100 mg/L; polymyxin B, 50 u/L; chloramphenicol, 10 mg/mL; streptomycin, 500 mg/L. X-Gal was used at 50 mg/L. Strains used in this work are listed in Supplementary Table 2.

### Strain construction

Primers used in this study are listed in Supplementary Table 3. We constructed chromosomal alterations in *V. cholerae* strains primarily using MuGENT (multiplex genome editing by natural transformation) (39), and also using allelic exchange with pKAS32 (40) as needed. Both methods have been previously described (5). The MuGENT method relies on natural transformation and co-transformation with a selectable marker at a neutral locus for in-frame deletions. We used *vc1807* as our neutral locus.

### Transposon mutagenesis screen

We mutagenized the Δ*flgC* Δ*vpsL* HCD-locked *vc1807*::Cm^R^ strain with Tn*5* as previously described (41). Mutants were selected on LB agar containing polymyxin B, kanamycin, and 0.5% glycerol. The addition of glycerol amplified differences in colony morphologies (42). Plates were incubated at 30^◦^C for 24 h and subsequently transferred to room temperature for 2 days. Exconjugants exhibiting reductions in opacity were isolated. Mutant strains were grown under aggregate-forming-conditions and screened for the loss of void formation at T = 22 h. Transposon insertion sites were determined using arbitrary PCR (43) and subsequently validated with primers specific to the identified loci.

### Aggregate formation

We used previously described aggregate forming conditions (5). To image aggregates, a 150 µL sample was placed into a well of a 1 No. 1.5 coverslip 96-well microtiter dish (MatTek, #P96G-1.5–5 F). To image voids, a 10 µL sample was gently mixed with 5 µL of India ink (Higgins, #44201) on a 60×44 mm coverslip (No. 1.5) and then a 44×44 mm coverslip (No. 1.5) was placed on top. Brightfield imaging was used to visualize and quantify void formation. Fluorescence imaging, as previously described (5), was used to visualize aggregate formation.

### Conditioned medium preparation

Strains were grown (10 mL total volume in 5 technical replicates) under aggregate forming conditions until T = 18 h and then pooled. Cultures were subjected to centrifugation for 30 min at 10,000 rpm on in a Sorvall RC 5B Plus centrifuge and then filter sterilized (pore size: 0.22 µm; MilliporeSigma, #SLGP033R). 2 mL of conditioned medium was placed into a test tube and returned to a rolling drum at 30° C until the designated time point. Samples were visualized as described above.

### Proteolytic zymography activity

Strains were grown overnight in LB medium with 0.1% arabinose at 37^◦^C with shaking (250 rpm). The optical densities (OD_600_) of cultures were measured and the cells were removed from the suspensions by centrifugation at 13,000 rpm for 10 min. The conditioned media were retained, sterilized by passage through a 0.22 µM filter (MilliporeSigma), the amounts normalized to the culture OD_600_ using water, and then concentrated ∼40-fold by passage over 10 kDa molecular cutoff spin columns (MilliporeSigma, #UFC901024 and #UFC501096). Proteins present in a 2 µL aliquot of each preparation were separated on a 10% SDS gel containing 0.1% gelatin (ThermoFisher, #ZY00105BOX) under non-reducing, non-denaturing conditions. The gel was washed twice with water for 5 min followed by incubation with renaturation buffer (ThermoFisher, #LC2670) for 21 h at 30° C. The gel was stained with Commassie Brilliant Blue R-250 (Biorad, #1610436) and imaged using trans-illumination. The look-up table was inverted using ImageJ (NIH). Thus, the presence of a band in the figure represents protease activity in that region of the gel.

### Proteolysis inhibition

Chemical inhibition of proteolytic activity was accomplished by dissolving a protease inhibitor cocktail tablet (cOmplete, Mini, EDTA-free; Roche, #04 693 159 001) into either 20 mL (high protease inhibitor concentration) or 100 mL (low protease inhibitor concentration) of LB medium containing 10 mM CaCl_2_. This medium was used to grow cultures under aggregate forming conditions.

### Immunoblotting

Conditioned media were obtained as described above. We combined 1 µL of these preparations with 4X SDS-PAGE buffer, boiled the samples for 20 min, and separated them on 4 to 20% Mini-Protein TGX gels (Bio-Rad, #4561096) at 100 V until the dye front reached the bottom of the gel (∼2 h). Proteins were transferred to polyvinylidene difluoride membranes (Bio-Rad, #1620174) for 1 h at 4°C and at 100 V. Membranes were incubated at room temperature for 40 min with a 1:5,000 dilution of monoclonal anti-FLAG-peroxidase antibody in PBST supplemented with 5% milk (Sigma, #A8592). The membranes were subsequently washed another 5 times with PBST. FLAG epitope-tagged protein levels were visualized using the Amersham ECL Western blotting detection reagent (GE Healthcare, #GERPN2209). In a parallel gel, 10 µL of the samples were separated on 4-20% Stain-Free TGX gels (Bio-Rad, #4568095), and assessed for total protein following the manufacturer’s recommendations. In all cases, samples were imaged (GE, ImageQuant, LAS 4000) and protein levels were quantified using ImageJ.

### Microscopy and image analysis

Microscopy and image analyses were performed as previously described (5). In brief, we used a Leica SP-8 point scanning confocal microscope equipped with a white-light laser for all confocal imaging. Samples were imaged using either a 10x air or 63x water objective. All features of aggregation and void formation were assessed using the 10x air objective. To determine onset timing of aggregation and void formation 1550×1550 µm^2^ regions were imaged every 1 h starting well before any perceivable void formation or aggregation had occurred. Samples were visually scored for the presence or absence of voids or aggregates. Onset timing for Void^+^ and Aggregate^+^ strains were assessed using the void-formation and aggregation-formation protocols, respectively. To quantify the sizes of voids and aggregates, samples were prepared using the void-formation protocol and three non-overlapping 4650×4650 µm^2^ regions were imaged using the tile scanning module. An intensity-based threshold segmentation on the brightfield channel was used to identify the extent of voids and aggregates across these regions. To determine the fractional occupancy of cells, an intensity-based segmentation was subsequently employed using the relevant fluorescent channel. All image analyses were performed in MATLAB using custom software (https://github.com/jemielita/aggregation.git). To visualize void formation and aggregation in strains lacking fluorescent reporters, SYTO-9 (ThermoFisher, #S34854; final concentration: 2.2 μM) was used. Cultures were first aliquoted into wells of microtiter dishes, dye was added, and samples were gently mixed.

### Growth rate analyses

Strains were grown as described above and 150 µL were added to wells of Corning 96-well Clear Flat Bottom Polystyrene TC-treated Microplates (Corning, #3598). Plates were incubated with shaking at 30^◦^C in a BioTek Synergy Neo2 Multi-Mode reader. OD_600_ was measured every 20 min for 16 h. Growth curves were normalized to the time when the cultures entered exponential phase, which we accomplished by shifting curves by the time at which the OD_600_ exceeded twice the minimum measured values.

## Acknowledgments

We thank Jon Paczkowski for help identifying the catalytic triad of Vca0812 and all members of the Bassler lab for insightful discussions. B.L.B. was supported by the Howard Hughes Medical Institute, National Science Foundation Grants MCB-1713731, NIH Grant 2R37GM065859, and the Max Planck Society-Alexander von Humboldt Foundation. M.L.J. was funded by a K99 Career Transition Award (1K99GM138764-01). A.A.M. is a Howard Hughes Medical Institute Fellow of the Life Sciences Research Institute. N.S.W. was supported by NIH funding (R01 GM082938). The funders had no role in study design, data collection and interpretation, or the decision to submit the work for publication.

---

**Supplementary Figure 1:**
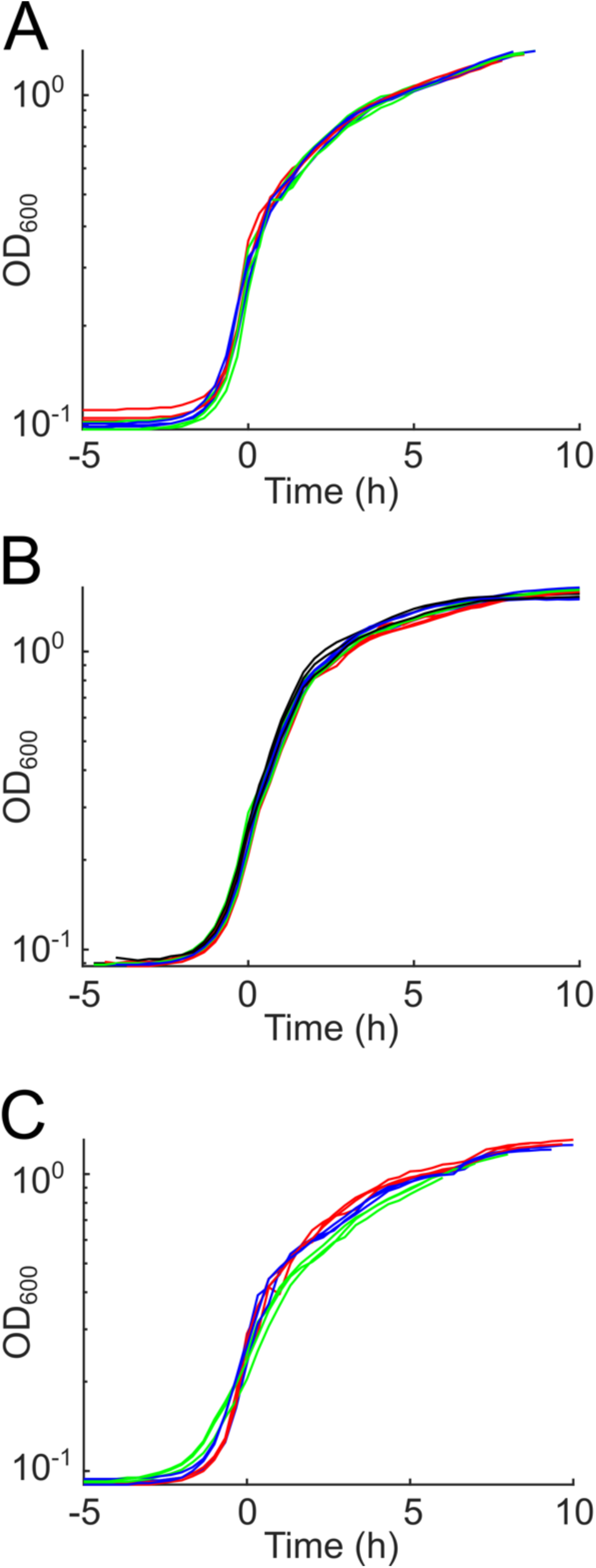
Growth curves for strains used in this work. (A) Growth curves for the Aggregate^+^ (red), Δ*vca0812* Aggregate^+^ (green), and Δ*vca0813* Aggregate^+^ (blue) strains. (B) Growth curves for Aggregate^+^ (red), Δ*hapA* Aggregate^+^ (green), Δ*prtV* Aggregate^+^ (blue), and Δ*4* Aggregate^+^ (black) strains. (C) Growth curves for the Aggregate^+^ strain to which no (red), a low (blue), or a high (green) concentration of a protease inhibitor cocktail was added at T = 0 h. (A-C) N = 3 biological replicates. All strains constitutively expressed *mKO* from the chromosome. Individual data traces were offset by the time at which the samples entered exponential growth.

**Supplementary Figure 2:**
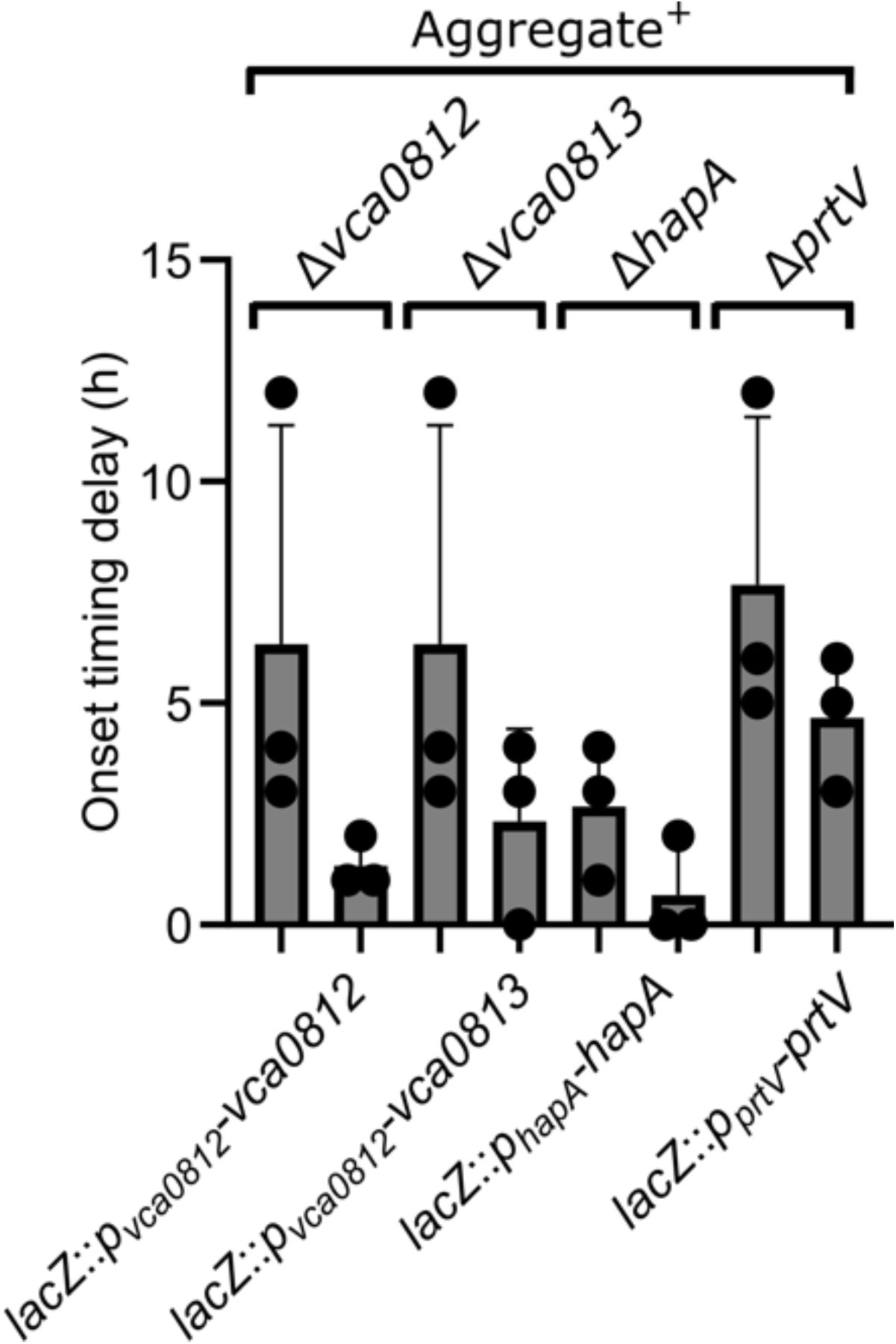
Protease complementation reduced aggregation onset timing delay. Quantification of aggregation onset timing delay for the designated strains relative to that of the Aggregate^+^ strain. Error bars denote mean ± SD, N = 3 biological replicates. Strains were visualized using the fluorescent stain SYTO-9.

**Supplementary Figure 3:**
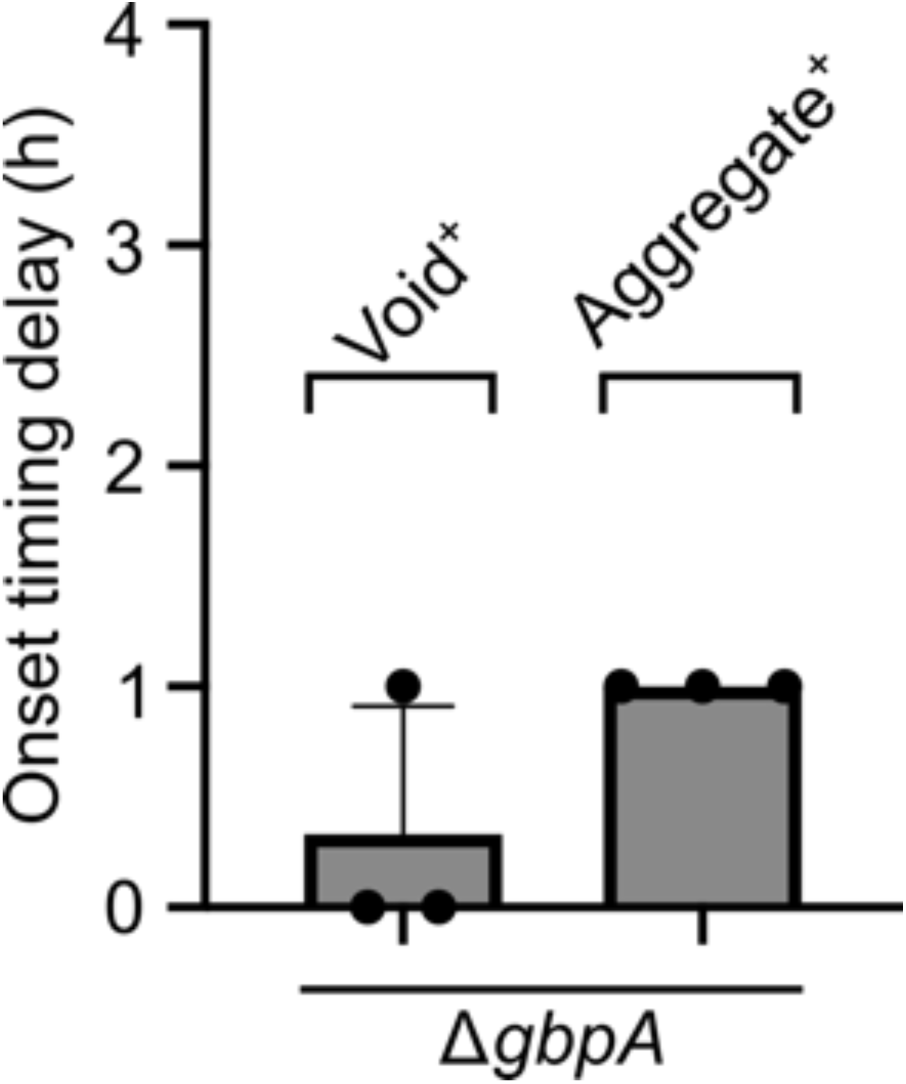
GbpA does not contribute to void formation or aggregation onset timing. Quantitation of void formation and aggregation onset timing delay for the designated strains relative to, respectively, the Void^+^ and Aggregate^+^ strains. Error bars denote mean ± SD, N = 3 biological replicates. All strains constitutively expressed *mKO* from the chromosome.

**Supplementary Figure 4:**
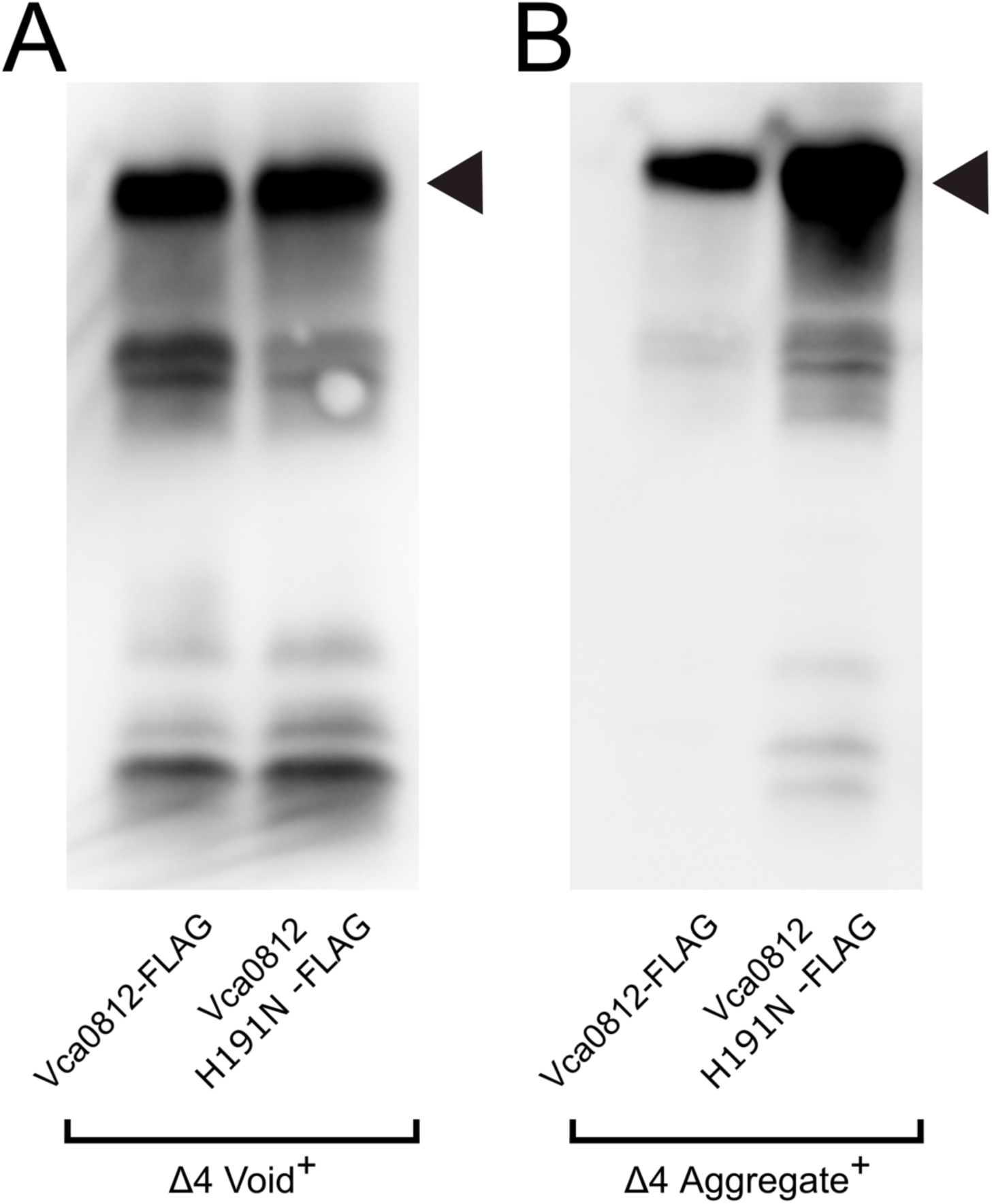
The Vca0812 and Vca0812 H191N proteins are produced at similar levels. Western blots showing the levels of the Vca0812-FLAG and Vca0812 H191N-FLAG proteins produced by the (A) Δ*4* Void^+^ strain and (B) Δ*4* Aggregate^+^ strain. The black arrows designate the positions of the Vca0812-FLAG and Vca0812 H191N-FLAG proteins.

**Supplementary Figure 5:**
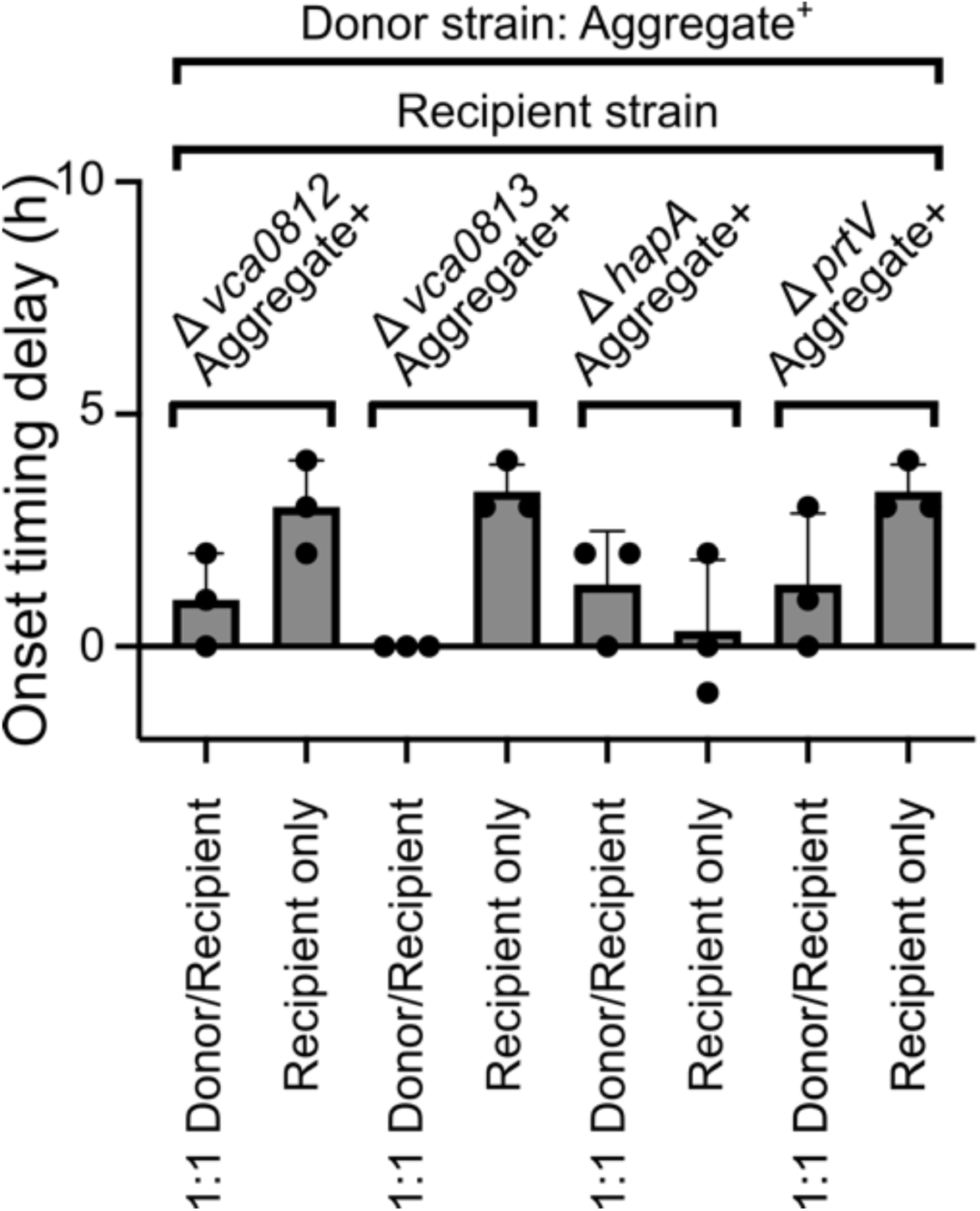
Co-culture of protease-deficient strains with the Aggregate^+^ strain restored aggregation onset timing. Quantitation of aggregation onset timing delay of the designated strains relative to that of the Aggregate^+^ strain. The donor strain is the Aggregate^+^ strain. Equal amounts of the donor and recipient were combined at T = 0 h. Error bars denote mean ± SD, N = 3 biological replicates. Donor and recipient strains, respectively, expressed *mKate2* and *mKO* from the chromosome.

**<u>Supplementary Video 1: Cross-feeding between Aggregate^+^ and Δ*4* Aggregate^+^ strains resulted in the formation of mixed-strain aggregates.</u>**

Representative *z*-scan at T = 24 h of a culture containing a 1:1 (v/v) mixture of the Aggregate^+^ and Δ*4* Aggregate^+^ strains. The Aggregate^+^ and Δ*4* Aggregate^+^ strains, respectively, constitutively expressed *mKate2* (red) and *mKO* (yellow) from the chromosome. Scale bar: 50 µm. Magnification: 63x.

**Supplementary Table 1:**
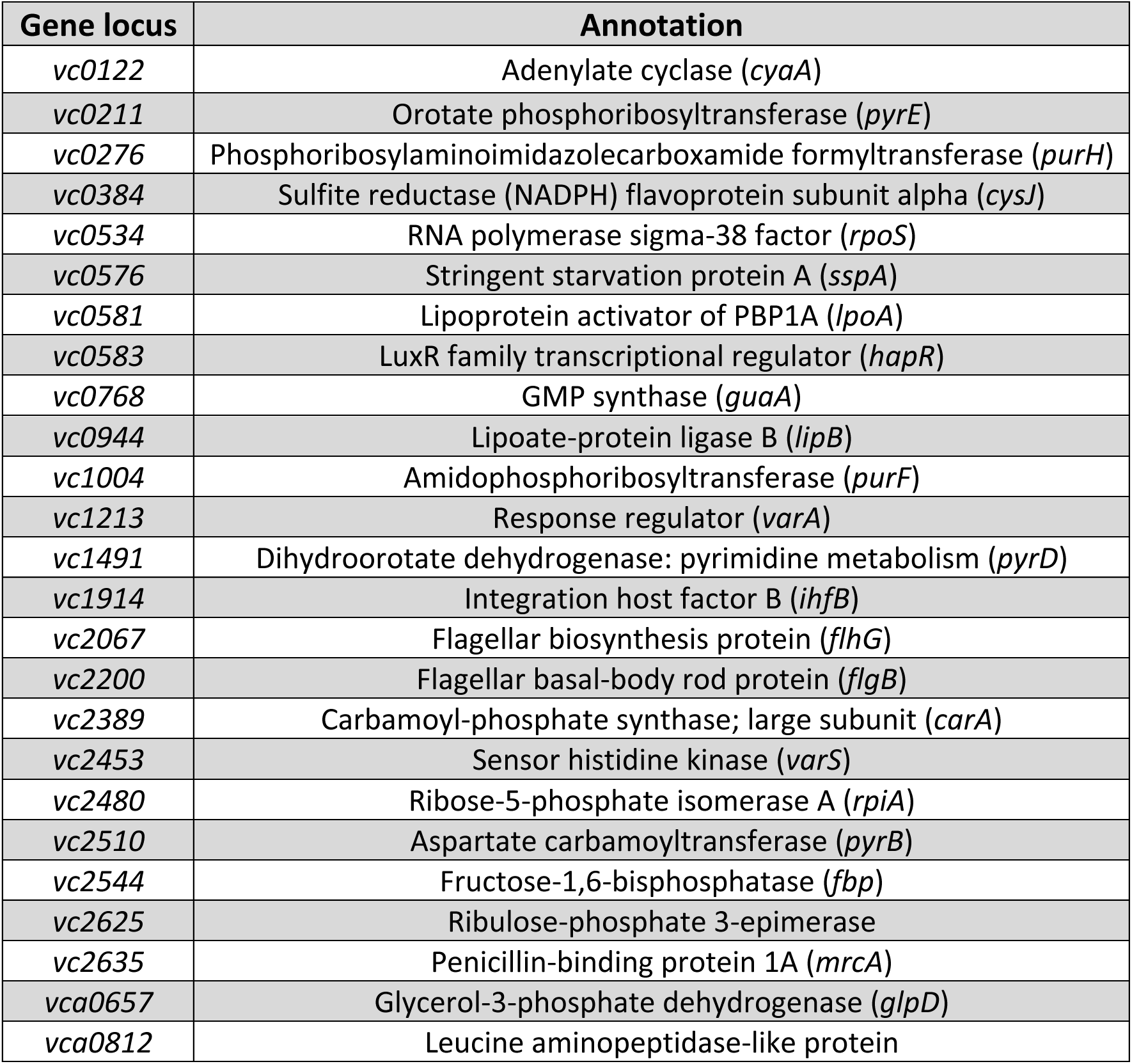
Genes identified in screen.

**Supplementary Table 2:**
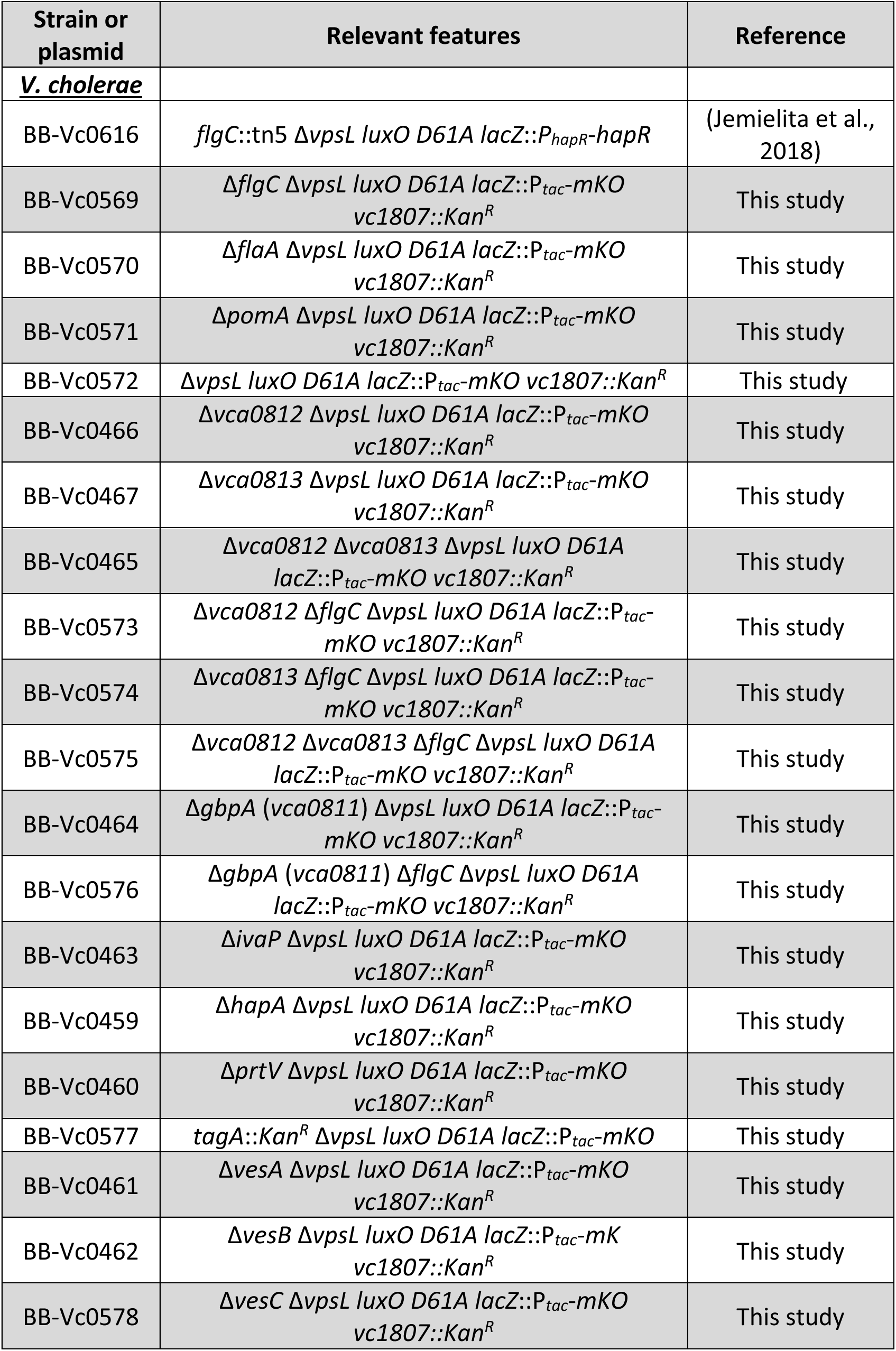
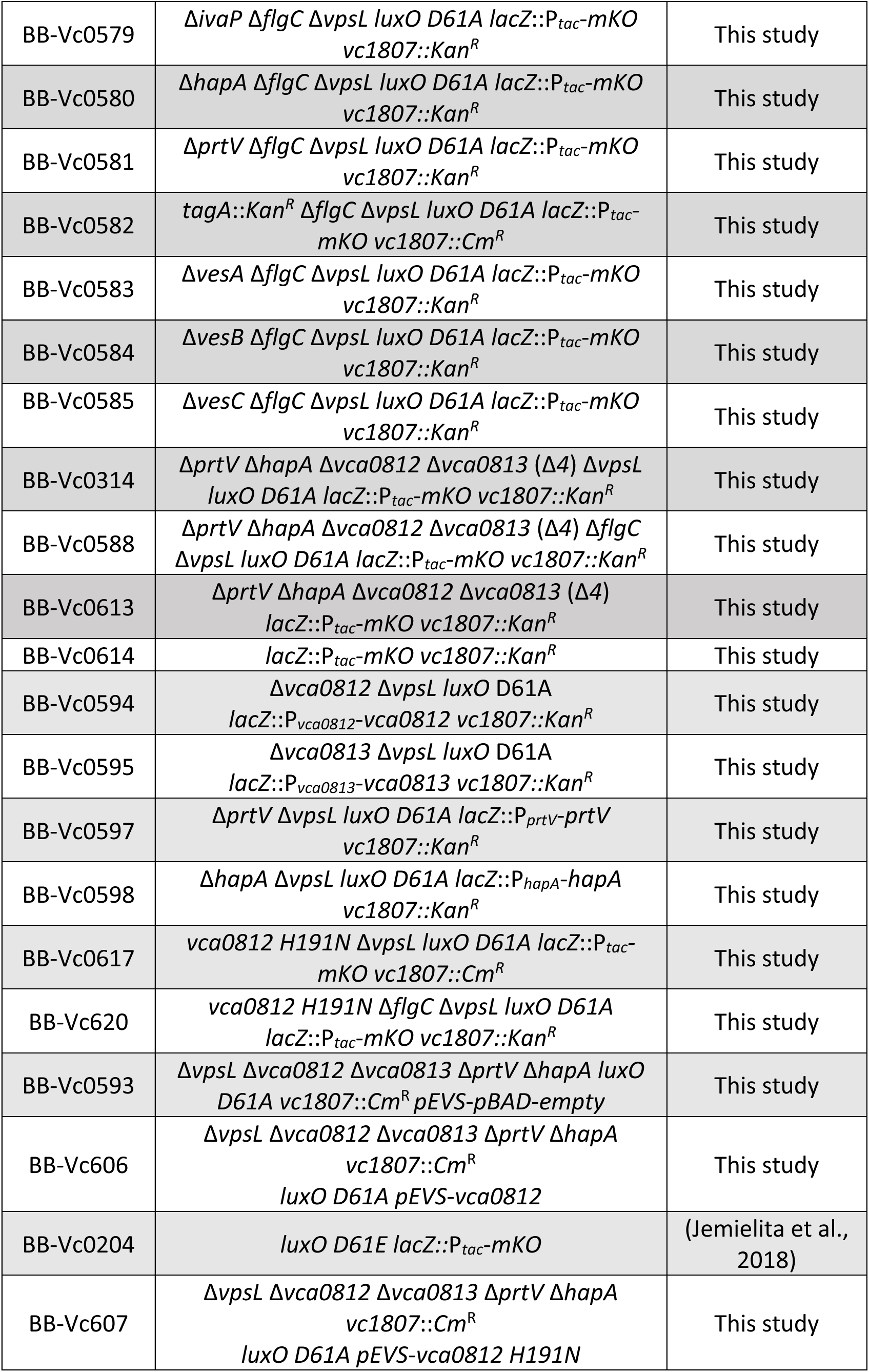
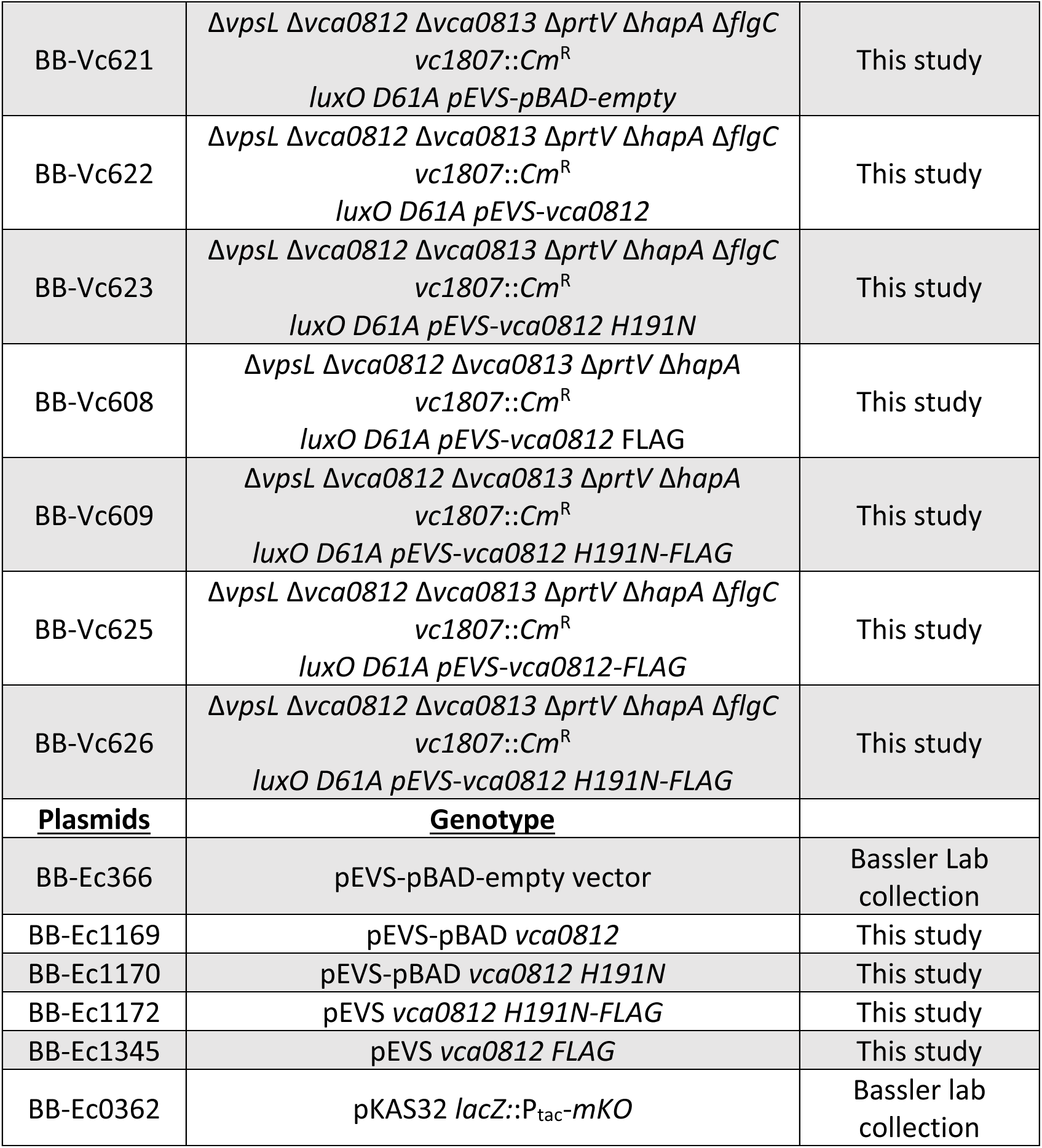
Strain list.

**Supplementary Table 3:**
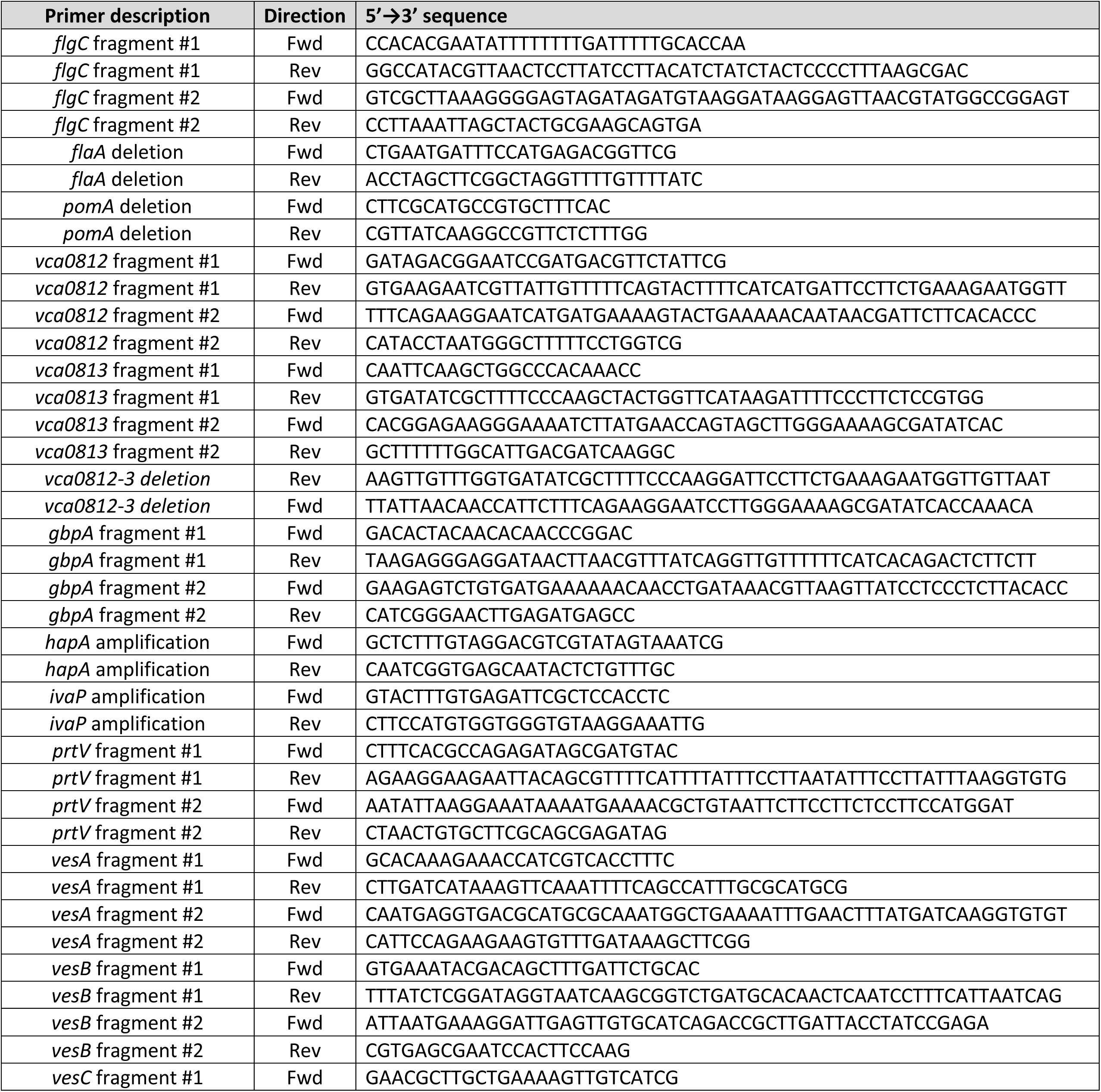
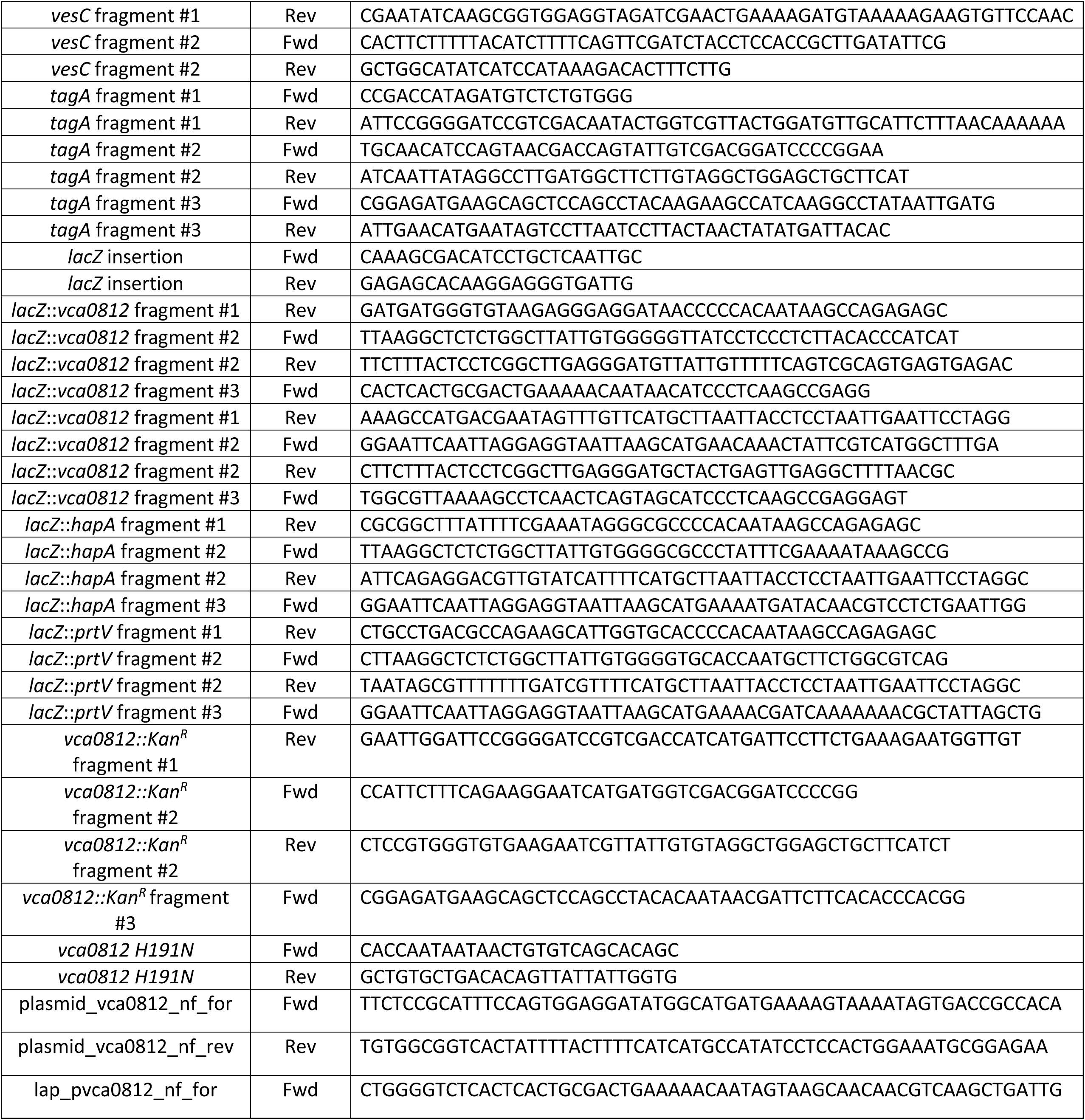
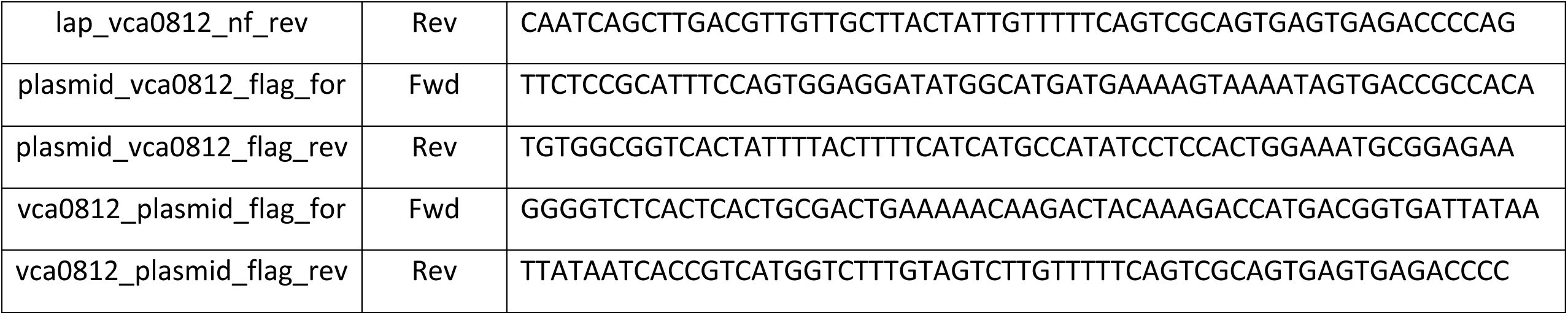
Primer list.

